# Sequential interactions with Mre11-Rad50-Nbs1 activate ATM/Tel1 at DNA double-strand breaks and telomeres

**DOI:** 10.1101/157305

**Authors:** Oliver Limbo, Yoshiki Yamada, Paul Russell

**Affiliations:** Department of Molecular Medicine, The Scripps Research Institute, La Jolla, CA 92037

**Keywords:** DNA damage signaling, DNA repair, DNA damage checkpoint, double strand break repair, telomeres

## Abstract

The Mre11-Rad50-Nbs1 (MRN) protein complex, CtIP/Ctp1/Sae2 and ATM/Tel1 kinase protect genome integrity through their functions in DNA double-strand break (DSB) repair, checkpoint signaling, and telomere maintenance. Nbs1 has a conserved C-terminal motif that binds ATM, but the full extent of ATM interactions with MRN are unknown. Here, we show that Tel1 overexpression in *Schizosaccharomyces pombe* restores Tel1 activity at DSBs and telomeres in the absence of Nbs1. This activity requires Mre11, indicating that Tel1 overexpression drives low affinity binding to the Mre11-Rad50 subcomplex. Mre11-Rad50 binds DSBs in *nbs1*Δ cells, and fusing the Tel1-binding motif of Nbs1 to Mre11 fully restores Tel1 signaling in these cells. Tel1 overexpression does not restore Tel1 signaling in cells carrying the *rad50-I1192W* mutation, which impairs the ability of Mre11-Rad50 to form the ATP-bound closed conformation. From these findings, we propose that Tel1 activation at DNA ends proceeds by a sequential mechanism initiated by high affinity binding to Nbs1 which recruits Tel1, followed by a low affinity interaction with Mre11-Rad50 in the closed conformation to activate Tel1.

## Introduction

DNA Double-strand breaks (DSBs) are one of the most dangerous lesions, as they break chromosomes as well as DNA. An inability to properly repair DSBs can result in cell death or cancer (Hoeijmakers, 2001). The Mre11-Rad50-Nbs1 (MRN) protein complex acts as a primary responder to DSBs, quickly localizing to damage sites (Stracker & Petrini, 2011). The FHA domain at the N-terminus of the Nbs1 subunit recruits CtIP/Ctp1, a DNA repair ortholog of *Saccharomyces cerevisiae* Sae2, which activates the intrinsic nuclease activity of Mre11 (Cannavo & Cejka, 2014, Limbo et al., 2007, Lloyd et al., 2009, Sartori et al., 2007, Williams et al., 2009). MRN-Ctp1 first nucleolytically displaces Ku or other proteins from DNA ends, and then initiates resection of the 5’ strand to generate a 3’ single-stranded DNA (ssDNA) overhang (Garcia et al., 2011, Lafrance-Vanasse et al., 2015, Langerak et al., 2011, Shibata et al., 2014). Rad51 mediates the invasion and base pairing of the ssDNA tail into homologous DNA sequences, usually in the sister chromatid, which it uses as a template to carry out the error-free pathway of homology-directed repair (Stracker & Petrini, 2011).

In addition to CtIP/Ctp1, the MRN complex also recruits the ATM/Tel1 (Ataxia telangiectasia mutated) serine/threonine kinase to damage sites (Uziel et al., 2003). ATM is a member of the PI3K-like protein kinase (PIKK) family of proteins, whose members include DNA-PKcs, ATR, and mTor (Paull, 2015). In response to damage, ATM serves to initiate cell cycle arrest, stimulate repair factors, and activate senescence and apoptosis pathways through the phosphorylation of many substrates that include CHK2, H2AX, NBS1, BRCA1, and p53 (Paull, 2015). Mutations in the ATM gene are associated with ataxia telangiectasia (A-T or Louis-Bar syndrome). Patients with this neurodegenerative disorder present with ataxia, telangiectasia, sensitivity to ionizing radiation, immunodeficiency, and a predisposition to cancer (Shiloh, 1997). Mutations in Nbs1 are associated with Nijmegen breakage syndrome, an A-T-like syndrome that includes microcephaly (Carney et al., 1998). Mre11 mutations are associated with an ataxia-telangiectasia-like disorder (ATLD), which resembles A-T with the exception that in most cases, ATLD does not result in cancer or immunodeficiency (Delia et al., 2004, Fernet et al., 2005, Stewart et al., 1999, Uchisaka et al., 2009).

The activities of the MRN complex are regulated by the Rad50 subunit, which provides a structural scaffold for the complex (Lafrance-Vanasse et al., 2015). Rad50 is an ABC-ATPase with an extended coiled-coil domain that is typical of SMC proteins. In the ATP-bound form, the Mre11-Rad50 globular domains, comprised of the Mre11 nuclease and Rad50 ATPase, are in a closed conformation promotes DNA binding/tethering and ATM activity. ATP hydrolysis leads to an opening of complex, which exposes the nuclease sites of Mre11 (Lammens et al., 2011, Lee et al., 2013, Lim et al., 2011, Mockel et al., 2012).

We have shown previously that a conserved domain in the C-terminus of the fission yeast and *Xenopus laevis* Nbs1 subunits binds ATM/Tel1 (You et al., 2005). This Tel1/ATM interaction mechanism is conserved in budding yeast Xrs2 and human Nbs1 (Falck et al., 2005). In human cells, elimination of the C-terminal 20-amino acids of Nbs1 impairs ATM localization into DNA repair foci and abrogates phosphorylation of some key substrates such as CHK2, (Falck et al., 2005). However, this motif is not required for ATM function in mice (Stracker et al., 2007). Purified human MRN was shown to stimulate ATM activity and DNA binding, but these also occur in the absence of Nbs1, albeit to a lesser extent (Lee & Paull, 2004, Lee & Paull, 2005). In *Xenopus* extracts, the Nbs1-ATM interaction is essential for both the recruitment of ATM to damage sites and its activity (You et al., 2005). Collectively, these data suggest that the C-terminal ATM/Tel1 binding motif of Nbs1 is highly conserved but there may be additional interaction mechanisms involving Nbs1 or even direct interactions of ATM/Tel1 with the Mre11-Rad50 subcomplex.

Here, we use fission yeast to probe for undiscovered interactions involving Tel1 and MRN *in vivo*. Our data suggests that high affinity binding to Nbs1 is principally responsible for localizing Tel1 at DSBs and telomeres, whereupon it is activated by a mechanism requiring the ATP-bound closed conformation of Mre11-Rad50.

## Results

### The C-terminus of Nbs1 is critical for Tel1 activity

The N-terminus of *S. pombe* Nbs1 contains an FHA domain fused to tandem BRCT domains. This region is flexibly linked to a C-terminal region containing Mre11 and ATM/Tel1 binding domains (Lafrance-Vanasse et al., 2015, Lloyd et al., 2009, Williams et al., 2009) (Fig 1A). These domains are conserved in mammalian Nbs1 proteins, but truncation of the C-terminal 24-amino acids in murine Nbs1 only partially impaired ATM activity (Stracker et al., 2007). This result might be explained if there is an additional MRN-ATM/Tel1 interaction mechanism. As *S. pombe* Nbs1 appears to have only a single Tel1-binding motif, which is located at the extreme C-terminus of the Nbs1 polypeptide, we decided to re-examine the importance of this motif for Te1l function using a sensitive immunoblot assay for formation of phosphorylated histone H2A, known as γH2A. As DNA is resected following DSB formation, the initial checkpoint signaling mediated by MRN-Tel1/ATM switches to Rad26/ATRIP-Rad3/ATR, correlating with the appearance of a 3’ ssDNA overhang (Limbo et al., 2011, Shiotani & Zou, 2009). Thus, both Rad3 and Tel1 contribute to phosphorylation of key substrates such as histone H2A, which is equivalent to H2AX in mammalian cells. Accordingly, we observed that ionizing radiation (IR)-induced γH2A formation is maintained in *tel1*Δ and *rad3*Δ single mutants but abolished in the double mutant (Fig 1B), as previously reported (Nakamura et al., 2004). To specifically assay the activity of Tel1, we performed subsequent assays in a *rad3*Δ background. γH2A formation was nearly abolished in *nbs1*Δ *rad3*Δ cells (Fig 1C), confirming the critical requirement of Nbs1 for Tel1 activity. Similarly, truncation of the C-terminal 60 residues of Nbs1 (*nbs1*-ΔC60) containing the Tel1-binding motif almost ablated γH2A formation specifically in the *rad3*Δ background (Fig 1C). The same defects were observed for double missense alleles *nbs1-9* (D603N, D604N) and *nbs1-10* (F611E, F613E) that mutate conserved residues in the Tel1 binding domain of Nbs1 (You et al., 2005). Interestingly, both *tel1Δ* and *nbs1-ΔC60* mutants had similar levels of γH2A (Fig 1B), underscoring the importance of the C-terminus of Nbs1 in Tel1 activity.

**Figure 1.**
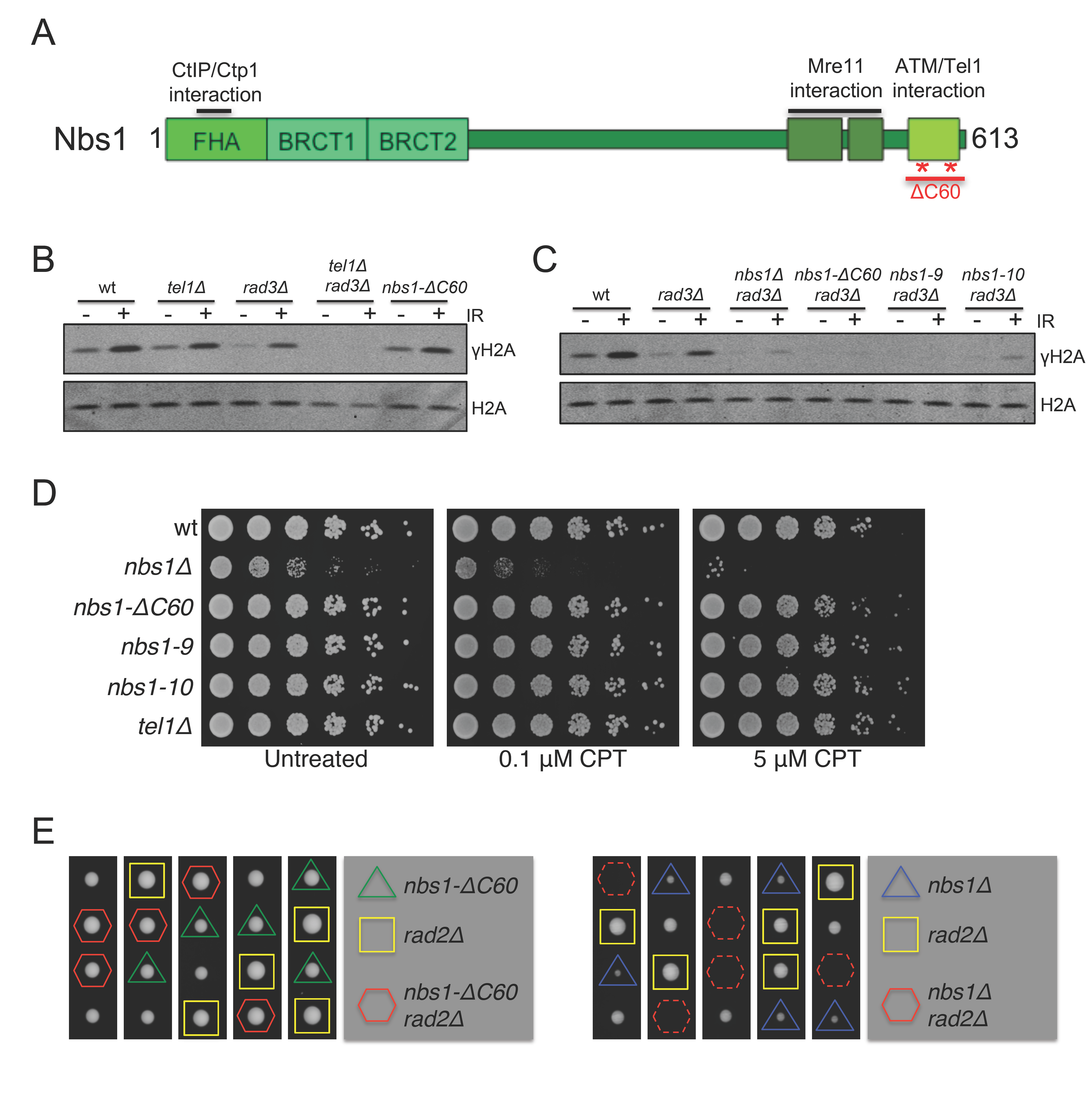
The C-terminus of Nbs1 is important for Tel1-mediated phosphorylation of histone H2A but dispensable for DNA repair. A) Domain architecture of Nbs1. Asterisks denote location of *nbs1-9* (D603N, D604N) and *nbs1-10* (F611E, F613E) alleles. B) Both Rad3 and Tel1 contribute to phosphorylation of histone H2A. Cells lacking the last 60 residues of Nbs1 encompassing the entire Tel1 binding domain (*nbs1-ΔC60*) have approximately the same level of γH2A as *tel1Δ*. C) *nbs1-ΔC60* and point mutations in the Tel1 interaction motif of Nbs1 *(nbs1-9*and *nbs1-10)* have greatly diminished Tel1-mediated γH2A formation. D) *Tel1Δ* and Nbs1 C-terminal mutants are insensitive to CPT. E) In the *rad2Δ*(FEN1) background,*nbs1-ΔC60*cells are viable (left),unlike *nbs1Δ*(right),as assayed by tetrad dissection.

In addition to its roles in Tel1 signaling, Nbs1 is also critical for DNA repair, recruiting the resection cofactor Ctp1 to damage sites through interactions with the N-terminal FHA domain (Lloyd et al., 2009, Williams et al., 2009). Indeed, cells lacking Nbs1 show severe growth defects and sensitivity to DNA damaging agents (Chahwan et al., 2003). To see if deletion or mutation of the Tel1 binding domain in the C-terminus affected the DNA damage repair function of Nbs1, we plated cells on plates containing the topoisomerase inhibitor, camptothecin (CPT). The sensitivity of *nbs1-ΔC60, nbs1-9* and *nbs1-10* mutants to CPT was not increased relative to wild-type (Fig 1D), as previously observed for IR and the alkylating agent, methyl methanesulfonate (MMS) (You et al., 2005). Similarly, *nbs1-ΔC60* cells were viable in the absence of the Rad2 (FEN1) flap endonuclease that is responsible for the maturation of Okazaki fragments, whereas *nbs1Δ* cells were lethal in this background (Fig 1E). Taken together, these data show that the C-terminal region of Nbs1 that is critical for Tel1 activity at DSBs is completely dispensable for the DNA repair functions of Nbs1.

### Overexpression of Tel1 bypasses the requirement for the C-terminus of Nbs1

Although the Tel1 interaction motif at the C-terminus of Nbs1 is critical for Tel1 activity at DSBs, we noted that in a *rad3*Δ background there was often a very weak but detectable IR-induced stimulation of γH2A in Nbs1 C-terminal mutants (Fig 1C). Surprisingly, we could also detect a weak IR-stimulated increase of γH2A in *nbs1*Δ *rad3*Δ cells. Importantly we never observed γH2A in *rad3*Δ *tel1*Δ cells (Fig 1B), confirming that Tel1 catalyzed the small amount of γH2A formed in *rad3*Δ *nbs1*Δ cells.

We suspected that the Nbs1-independent activity of Tel1 might involve a low affinity interaction of Tel1 with Mre11-Rad50 (MR). To address this possibility, we first investigated whether overexpression of Tel1 bypasses the requirement for the Tel1 binding domain of Nbs1. We transformed *nbs1-ΔC60 rad3Δ* cells with a plasmid expressing *tel1*^+^ driven from the attenuated thiamine-repressible *nmt1* promoter, or an empty vector control, and found that Tel1 overexpression restored both basal and IR-induced phosphorylation of H2A (Fig 2A).

**Figure 2.**
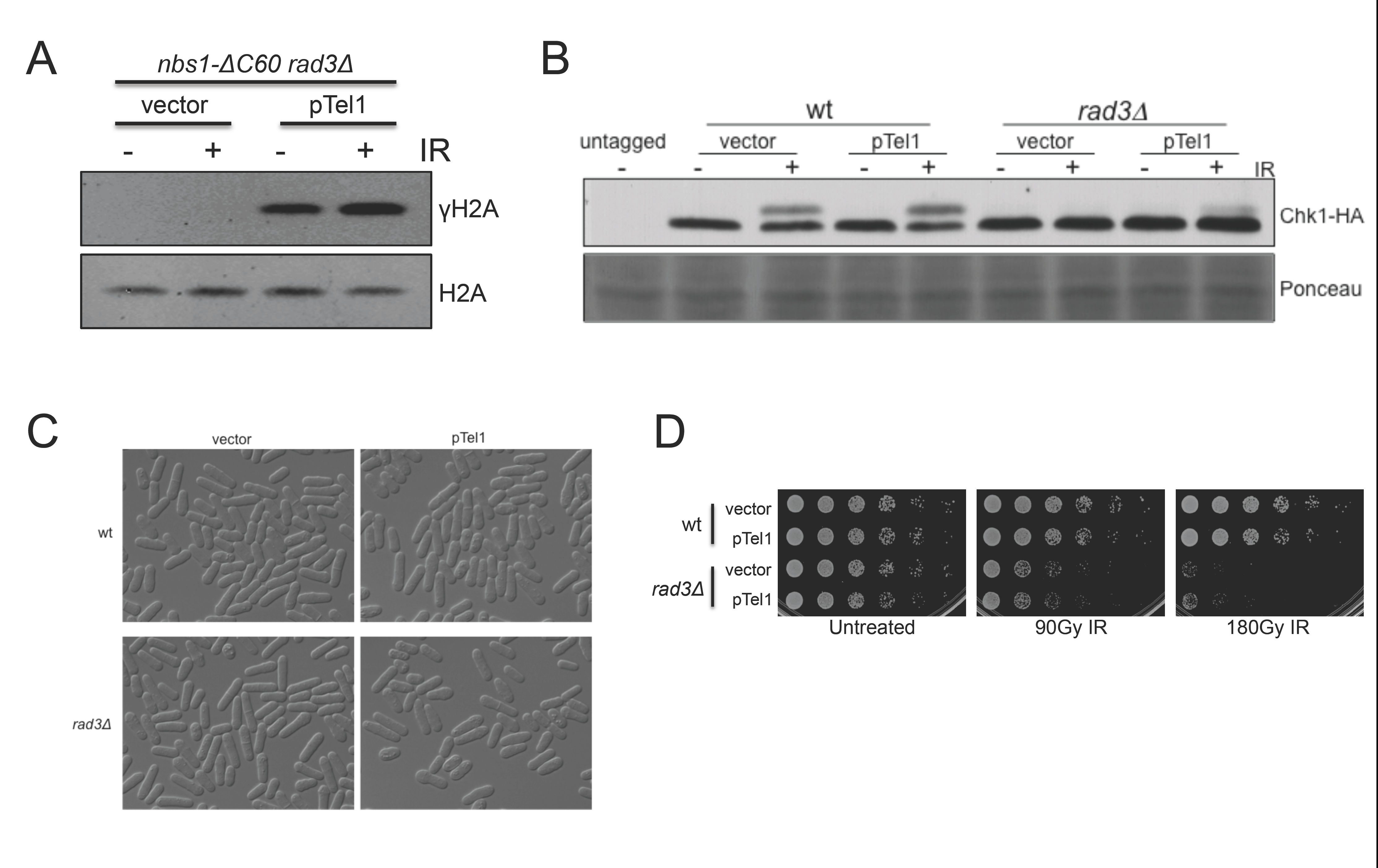
Overexpression of Tel1 bypasses the need for the C-terminus of Nbs1 in Tel1 signaling without deleterious effects. A) *nbs1-ΔC60 rad3Δ* cells were transformed with a plasmid containing *tel1+* expressed from the *nmt41* promoter, or an empty vector, and assayed for γH2A formation following IR exposure. B) Tel1 overexpression does not spontaneously cause activating phosphorylation of Chk1. However, Tel1 overexpression does partially restore Chk1 phosphorylation in *rad3Δ* cells in response to IR. C) Tel1 overexpression does not affect cell morphology or cause cell elongation. D) Tel1 overexpression does not negatively affect growth rate nor increase sensitivity to IR.

Overexpression of Tel1 in *S. cerevisiae* caused prolonged cell-cycle arrest via Rad53, even in the absence of exogenous DNA damage (Clerici et al., 2001). To address whether Tel1 overexpression alone triggered the DNA damage checkpoint in *S. pombe*, we first assayed phosphorylation of the effector checkpoint kinase, Chk1. In the absence of damage, Tel1 overexpression did not cause a detectable mobility shift in Chk1 in either wild type or *rad3Δ* backgrounds (Fig 2B). Rad3 mediates phosphorylation of Chk1 in most contexts (Limbo et al., 2011). Interestingly, overexpression of Tel1 partially restored IR-dependent Chk1 phosphorylation in *rad3Δ* cells (Fig 2B). Cells overexpressing Tel1 had normal cell morphology and were not hyper-elongated, indicating that Tel1 overexpression did not activate Cds1/Rad53 or Chk1 (Fig 2C). Moreover, Tel1 overexpression did not cause obvious growth defects nor increased sensitivity to DNA damaging agents (Fig 2D), suggesting that overexpression of Tel1 had no overt negative consequences to the cells.

### Tel1 overexpression bypasses the requirement of Nbs1, but not Mre11, in DNA damage-induced Tel1 activity and telomere maintenance

Having found that Tel1 overexpression can bypass the requirement for the C-terminus of Nbs1, we next examined whether the same was true in cells lacking Nbs1 or Mre11. In *nbs1Δ rad3Δ* cells, Te1l overexpression rescued both basal and IR-induced phosphorylation of histone H2A (Fig 3A). However, IR-induced formation of γH2A was defective in *mre11Δ rad3Δ* cells overexpressing Tel1, with only basal γH2A levels being restored (Fig 3A). Similarly, Te1l overexpression partially restored IR-induced Chk1 phosphorylation in *nbs1Δ rad3Δ* cells, but not *mre11Δ rad3Δ* cells (Fig 3B).

**Figure 3.**
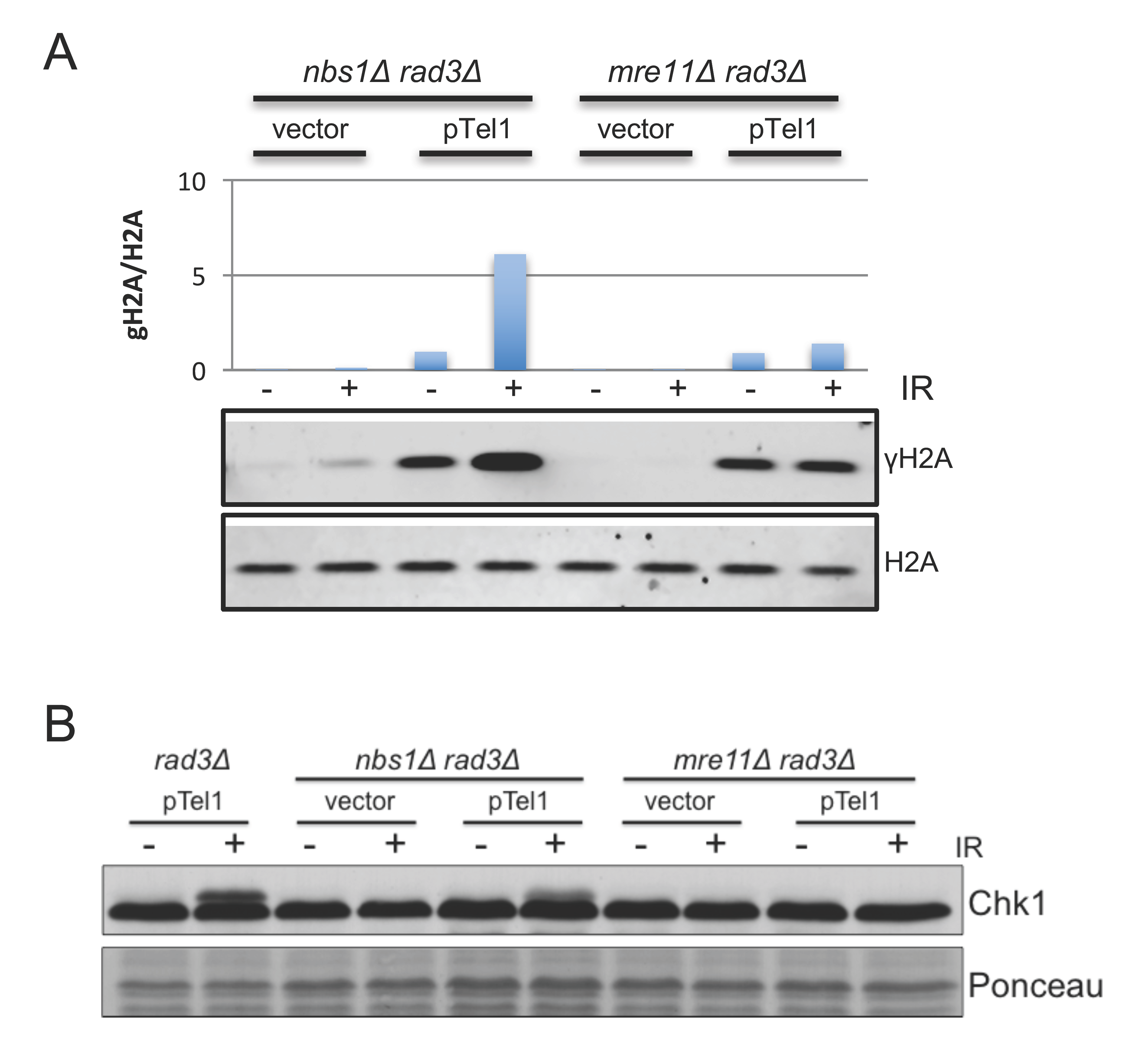
Overexpression of Tel1 bypasses the requirement of Nbs1, but not Mre11, in damage-induced Tel1 signaling. A) Tel1 overexpression in *nbs1Δ rad3Δ* restores basal and IR-induced γH2A formation. In the *mre11Δ rad3Δ* background, only basal levels of γH2A are restored. B) Chk1 phosphorylation in response to IR is partially restored when Tel1 is overexpressed in the *nbs1Δ rad3Δ* background but not in *mre11Δ rad3Δ* cells.

In *S. pombe*, both Rad3 and Tel1 contribute to maintenance of telomeres through phosphorylation of Ccq1, a subunit of the Shelterin complex. Phosphorylated Ccq1 then promotes recruitment of telomerase (Moser et al., 2011, Yamazaki et al., 2012). Thus, in the absence of Rad3 and Tel1, cells undergo telomere erosion with a small subset surviving through circularization of chromosomes after successive passages (Nakamura et al., 2002). A similar relationship was observed when MRN null mutants were combined with *rad3Δ* mutants, underscoring the importance of the MRN complex in Tel1 activity at telomeres. We asked whether Tel1 overexpression could prevent telomere loss in cells lacking Rad3 and subunits of MRN complex. To address this question, we generated *mre11Δ rad3Δ* or *nbs1Δ rad3Δ* mutants with *tel1*^+^ either driven from its native promoter or the *nmt1* overxpression promoter. Genomic DNA was prepared from cells after each passage and Southern blotting was performed probing for telomere-associated sequence (TAS1). We found overexpression of Tel1 prevented the loss of telomeres in *nbs1Δ rad3Δ* cells (Fig 4). Tel1 overexpression also had an effect in *mre11Δ rad3Δ* cells, although in this case it only delayed the loss of telomeres (Fig 4). These effects correlated with the improved growth of *nbs1Δ rad3Δ* and *mre11Δ rad3Δ* cells in the presence of overexpressed Tel1 (Fig EV1). Taken together, our results show that Tel1 overexpression can bypass the requirement for Nbs1, but not Mre11, in both DNA damage signaling and telomere maintenance.

**Figure 4.**
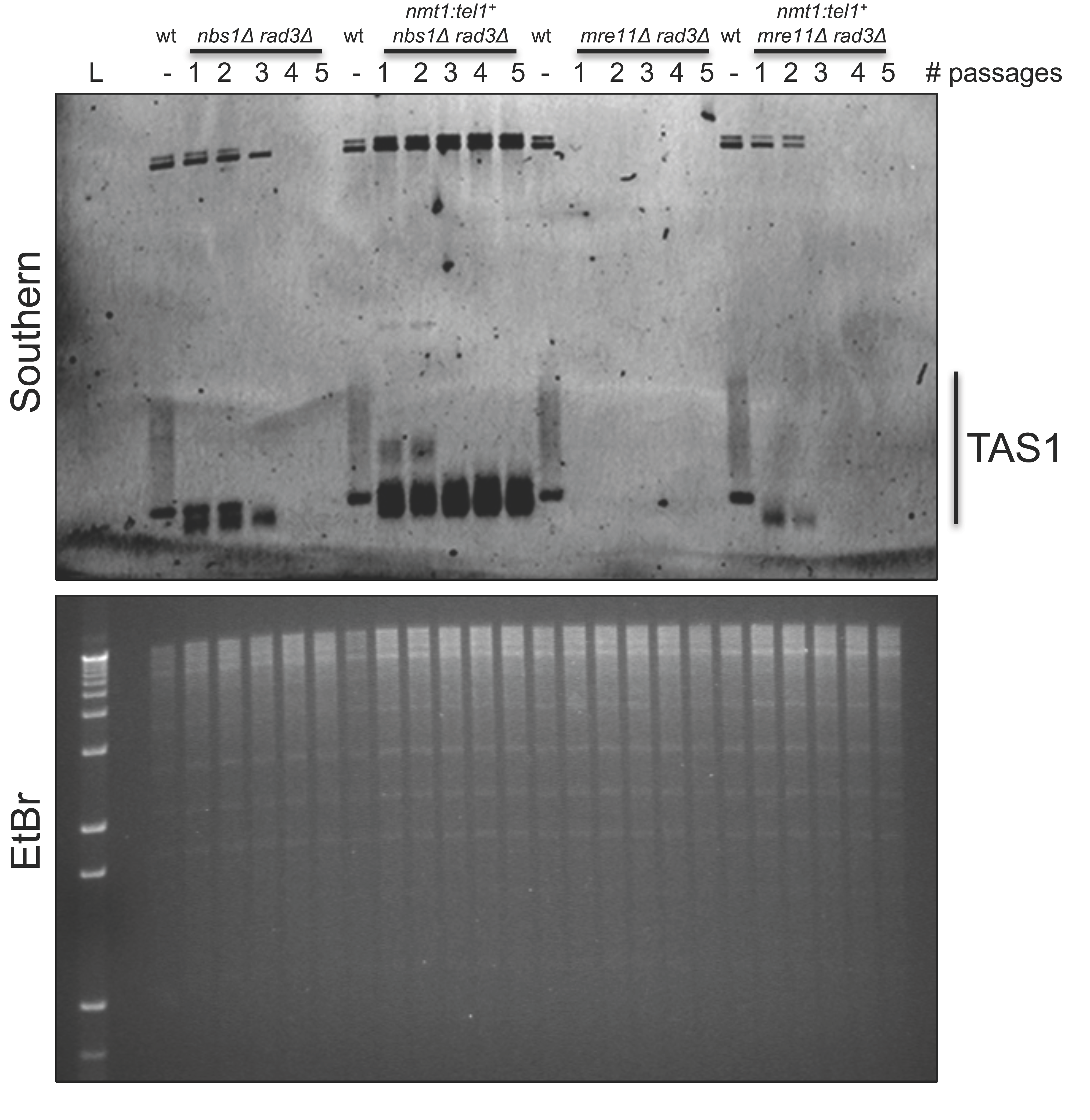
Overexpression of Tel1 bypasses the requirement of Nbs1, but not Mre11, in Tel1-mediated telomere maintenance. Southern blot of EcoRI digested genomic DNA probing for Telomere Associated Sequences (TAS1). Telomere loss observed in *nbs1Δ rad3Δ* cells after successive passages is prevented by Tel1 overexpression. In the *mre11Δ rad3Δ* background, Tel1 overexpression only slightly delayed telomere erosion. Ethidium bromide (EtBr) stained gel serves as a loading control.

### Mre11-Rad50 form a complex independently of Nbs1 that is capable of binding DSBs

The different consequences of Tel1 overexpression in *mre11Δ* and *nbs1Δ* backgrounds were surprising, given they are part of the same protein complex and both mutants are equally sensitive to DNA damaging agents (Chahwan et al., 2003). These results prompted us to examine whether Mre11 and Rad50 can form a stable protein complex without Nbs1. We performed co-immunoprecipitation experiments using strains with Mre11-MYC and TAP-Rad50 expressed from their endogenous loci and under their native promoters. Interestingly, Mre11-MYC co-precipitated readily with TAP-Rad50, both in the presence and absence of Nbs1 (Fig 5A).

**Figure 5.**
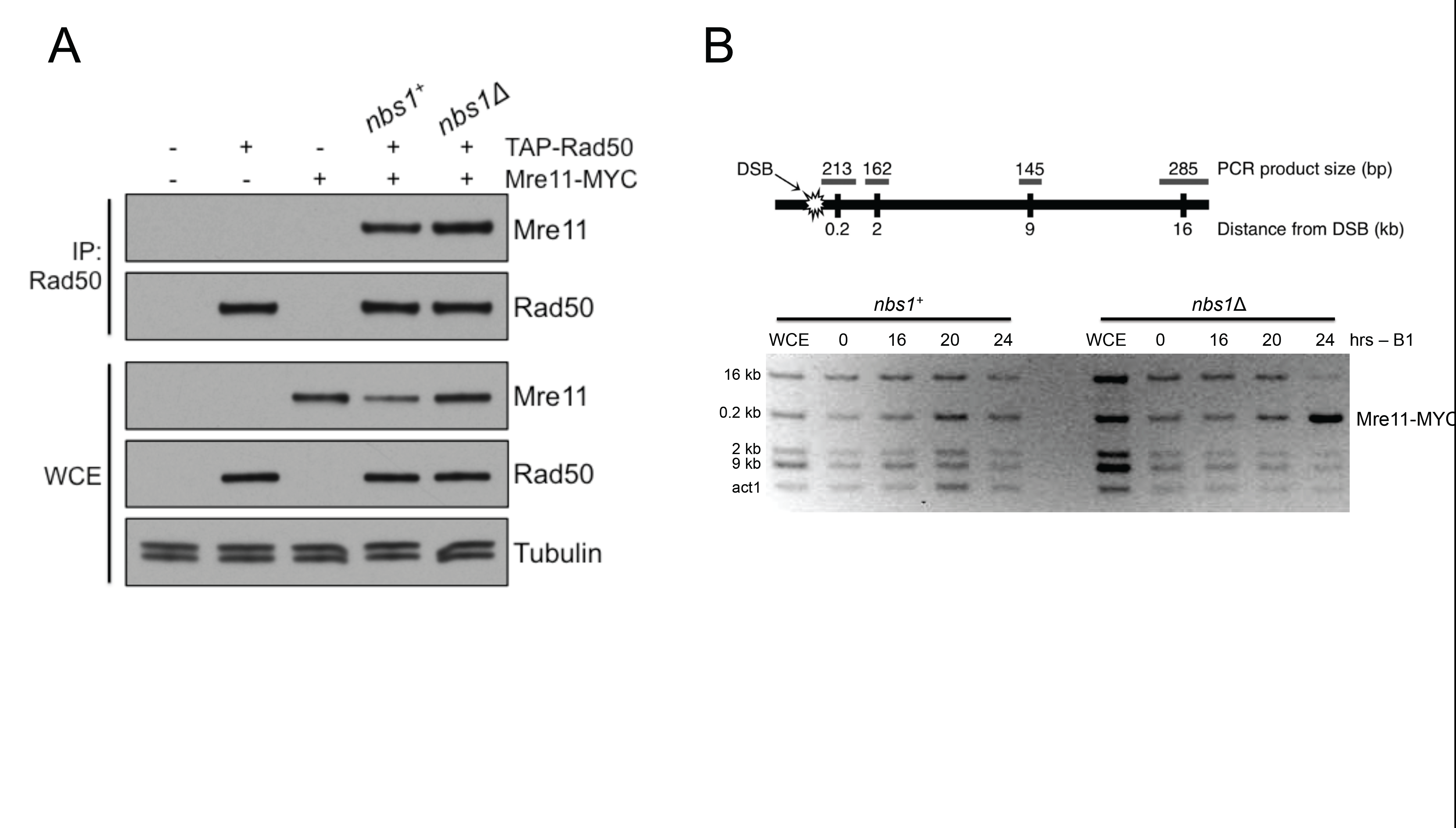
Mre11-Rad50 form a complex independently of Nbs1 that can localize to DNA double-strand breaks. A) Mre11-MYC efficiently co-precipitates with TAP-Rad50 in the presence or absence of Nbs1. B) Mre11-MYC is efficiently enriched at a DSB created by HO endonuclease with or without Nbs1 as assayed by chromatin immunoprecipitation. Note that derepression of HO endonuclease expression under the control of the *nmt1* promoter occurs at about 16-22 hours after removal of thiamine from the growth media (Forsburg & Rhind, 2006).

We next performed a chromatin immunoprecipitation (ChIP) experiment, assaying Mre11 enrichment around a site-specific DSB generated by the HO endonuclease. Mre11 was enriched immediately adjacent (0.2kb) from the break site in both wild-type and *nbs1Δ* backgrounds (Fig 5B). In fact, there was a greater enrichment of Mre11 in the *nbs1Δ* background, which we previously observed in *ctp1Δ* and Mre11 nuclease-deficient mutants, which are defective in resection but maintain the integrity of the MRN protein complex (Limbo et al., 2011, Williams et al., 2008). Taken together, our results indicate that Mre11-Rad50 can form a stable subcomplex in the absence of Nbs1 that can localize to DSBs.

### Fusion of the C-terminus of Nbs1 to Mre11 restores Tel1 activity in *nbs1Δ* cells

Our results suggested that Tel1 has a high-affinity interaction with the C-terminus of Nbs1, which can be bypassed by promoting a low affinity interaction with Mre11-Rad50 through Tel1 overexpression. However, our results did not exclude a model in which Nbs1 also has a role in activating Tel1 that is overcome by Tel1 overexpression. To address this question, we fused the last 60 residues of Nbs1 containing the Tel1 binding domain (TBD) to the C-terminus of full-length Mre11 (Fig 6A). This *mre11-TBD* construct replaced the endogenous *mre11*^+^ gene. We then assayed Tel1 activity in the *mre11-TBD nbs1Δ rad3Δ* background. The Mre11-TBD fusion fully restored phosphorylation of histone H2A to at least the level of *rad3Δ* cells, both basally and in response to IR treatment (Fig 6B). In fact, the basal level of γH2A exceeded that of *rad3Δ* cells. Moreover, we found that the Mre11-TBD fusion prevented the telomere erosion observed in *nbs1Δ rad3Δ* cells (Fig 6C).

**Figure 6.**
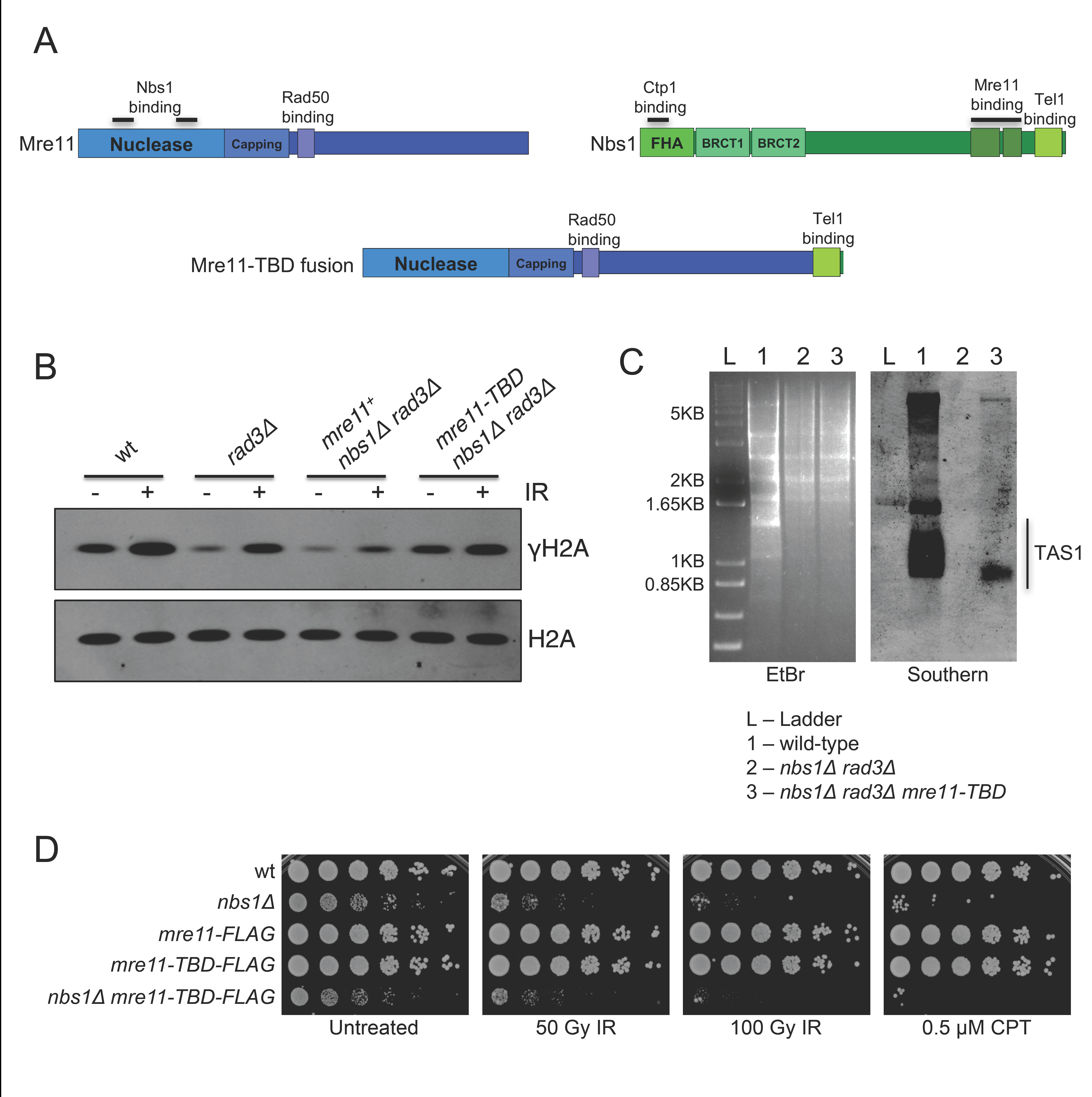
Fusion of the Nbs1 C-terminus to Mre11 is sufficient for Tel1 activity. A) Schematic of Mre11, Nbs1, and fusion protein generated by addition of C-terminal 60 amino acids encompassing the Tel1 binding domain of Nbs1 to the C-terminus of full-length Mre11. B) The Mre11-TBD (Tel1 Binding Domain) fusion protein restores histone H2A phosphorylation in the *nbs1Δ rad3Δ* background under endogenous Tel1 levels. C) The Mre11-TBD fusion protein prevents telomere loss in *nbs1Δ rad3Δ* cells under endogenous Tel1 levels. Strains were passaged 10 times and EcoRI-digested genomic DNA was probed for Telomere Associated Sequences (TAS1). Ethidium bromide (EtBr) stained gel serves as a loading control. D) The Mre11-TBD fusion protein does not noticeably affect Mre11 function in response to IR or CPT. The fusion protein does not correct the DNA repair defect of *nbs1Δ* cells.

To test the effect of the Mre11-TBD fusion on Mre11 function, we performed spot dilution assays, exposing the strains to different DNA damaging agents (Fig 6D). The fusion protein alone did not increase sensitivity of cells to IR and CPT, suggesting it did not impair Mre11 function. As expected, the Mre11-TBD fusion did not restore the DNA damage repair defect of *nbs1Δ* cells, because the DNA repair activity of MRN protein complex requires Ctp1 binding to the FHA domain found at the N-terminus of Nbs1. Taken together, these results show that fusion of the C-terminal 60 residues of Nbs1 to Mre11 was sufficient to restore Tel1 signaling in *nbs1Δ* cells, but the DNA repair defect remained.

### Nbs1-independent activity of Tel1 at DSBs requires ATP-bound closed conformation of Mre11-Rad50

Our results showed that the functions of Nbs1 in DNA damage-induced Tel1 activation could be largely bypassed by overexpression of Tel1 or fusion of the Tel1-binding domain of Nbs1 to Mre11. However, Tel1 overexpression had no effects in the absence of Mre11. Thus, it is likely that Mre11-Rad50, which can bind damage sites independently of Nbs1, has a low affinity interaction with Tel1 that is sufficient to restore Tel1 activity to *nbs1Δ* cells when Tel1 is overexpressed. To explore if this role of Mre11- Rad50 in Tel1 activity required Mre11 nuclease activity, we repeated our γH2A assay in cells with the *mre11-H134S* allele, which ablates Mre11 nuclease activity (Williams et al., 2008). Tel1 overexpression restored basal and IR-induced γH2A formation in *mre11-H134S nbs1Δ rad3Δ* cells (Fig 7A), indicating that the nuclease activity of Mre11 is not required for the MR-dependent activity of Tel1.

**Figure 7.**
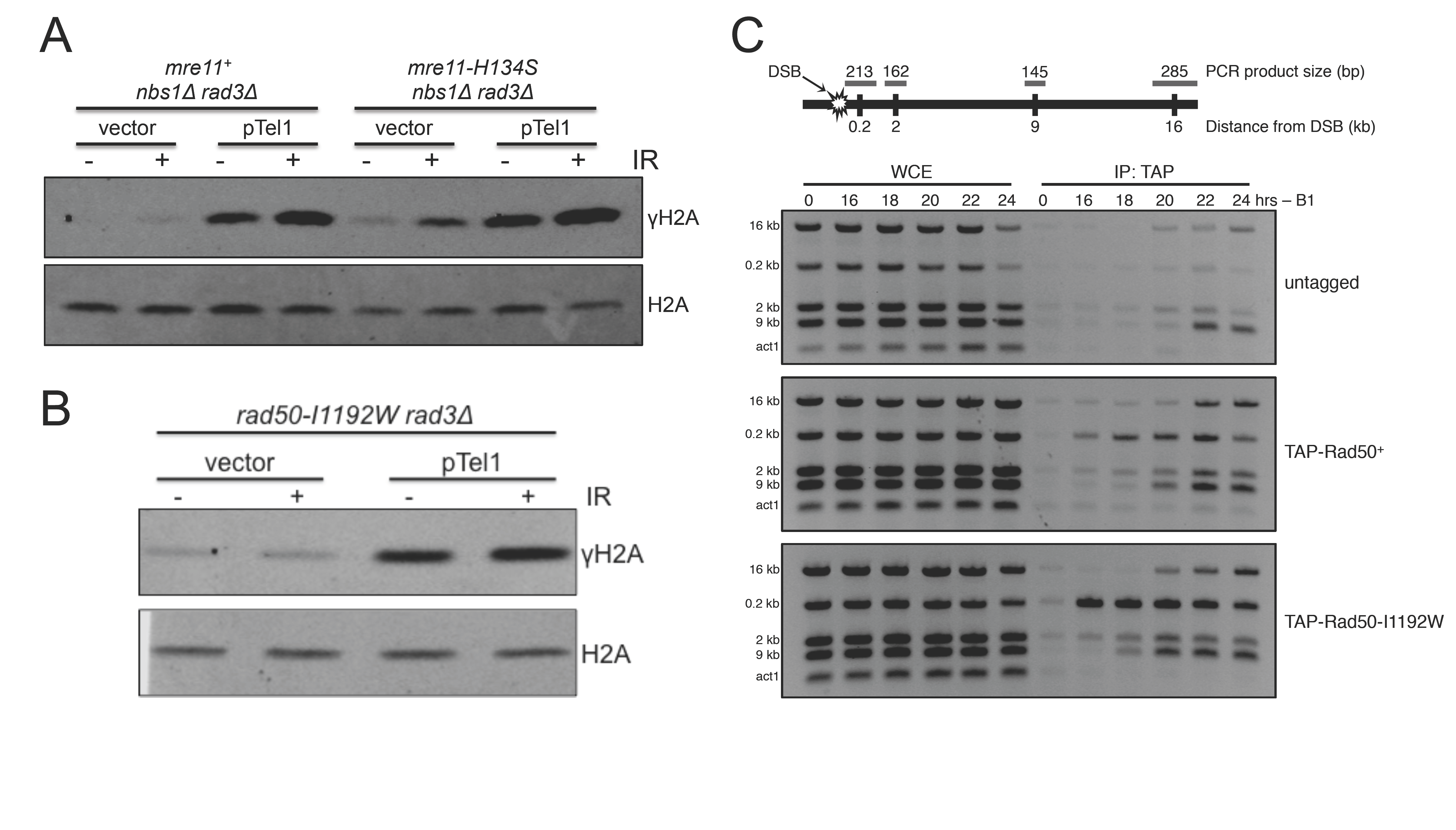
Stimulation of Tel1 activity by Mre11-Rad50 depends on conformational state but not nuclease activity. A) Nuclease-dead, *mre11-H134S*, does not reduce γH2A formation in *nbs1Δ rad3Δ* cells when Tel1 is overexpressed. B) The open-conformation *TAP-rad50-I1192W* allele is unable to stimulate Tel1 activity in response IR damage. C) TAP-Rad50-I1192W is enriched at DNA double-strand breaks induced by the HO-endonuclease.

Upon binding ATP, the Mre11-Rad50 subcomplex undergoes a conformational switch from an open to a closed state (Lammens et al., 2011, Lim et al., 2011, Mockel et al., 2012). The Rad50-I1192W mutation interferes with the conformational switch by obstructing a cavity in the dimer that accommodates the closed conformation. We previously showed that TAP-tagged Rad50-I1192W is partially defective in DSB repair and nearly completely defective in Tel1 signaling (Williams et al., 2011). Overexpression of Tel1 was unable to overcome the deficiency of the *TAP-rad50-I1192W rad3Δ* mutant to generate IR-induced γH2A (Fig 7B). Importantly, the mutant Rad50 protein was readily detected at an HO-induced DSB (Fig 7C), indicating that it maintained the ability to form a complex with Mre11 that binds DSBs. Thus, overexpression of Tel1 does not overcome the requirement for the proper conformation of MR complex in detecting Tel1 activity at DSBs.

## Discussion

In this study, we have uncovered evidence for a mechanism of sequential recruitment and activation of Tel1/ATM at DSBs and telomeres. In fission yeast, the Tel1-binding module at the C-terminus of Nbs1 is critical for Tel1 function, but this recruitment mechanism can be bypassed by increasing the cellular concentration of Tel1. Indeed, the entirety of Nbs1 protein is dispensable for Tel1 signaling when Tel1 is overexpressed. However, Mre11 remains essential for Tel1 activity at DSBs and telomeres, even when Tel1 is overexpressed. This result implies that Mre11-Rad50 localizes in the nucleus, binds DSBs, and maintains a low affinity interaction with Tel1 in the absence of Nbs1. Indeed, we detected strong enrichment of Mre11 at a DSB in *nbs1Δ* cells. From these results, we propose that for Tel1 activity at DSBs and telomeres, Nbs1 principally serves a recruitment or enrichment role, whereas Mre11-Rad50 plays a critical stimulatory role (Fig 8).

**Figure 8.**
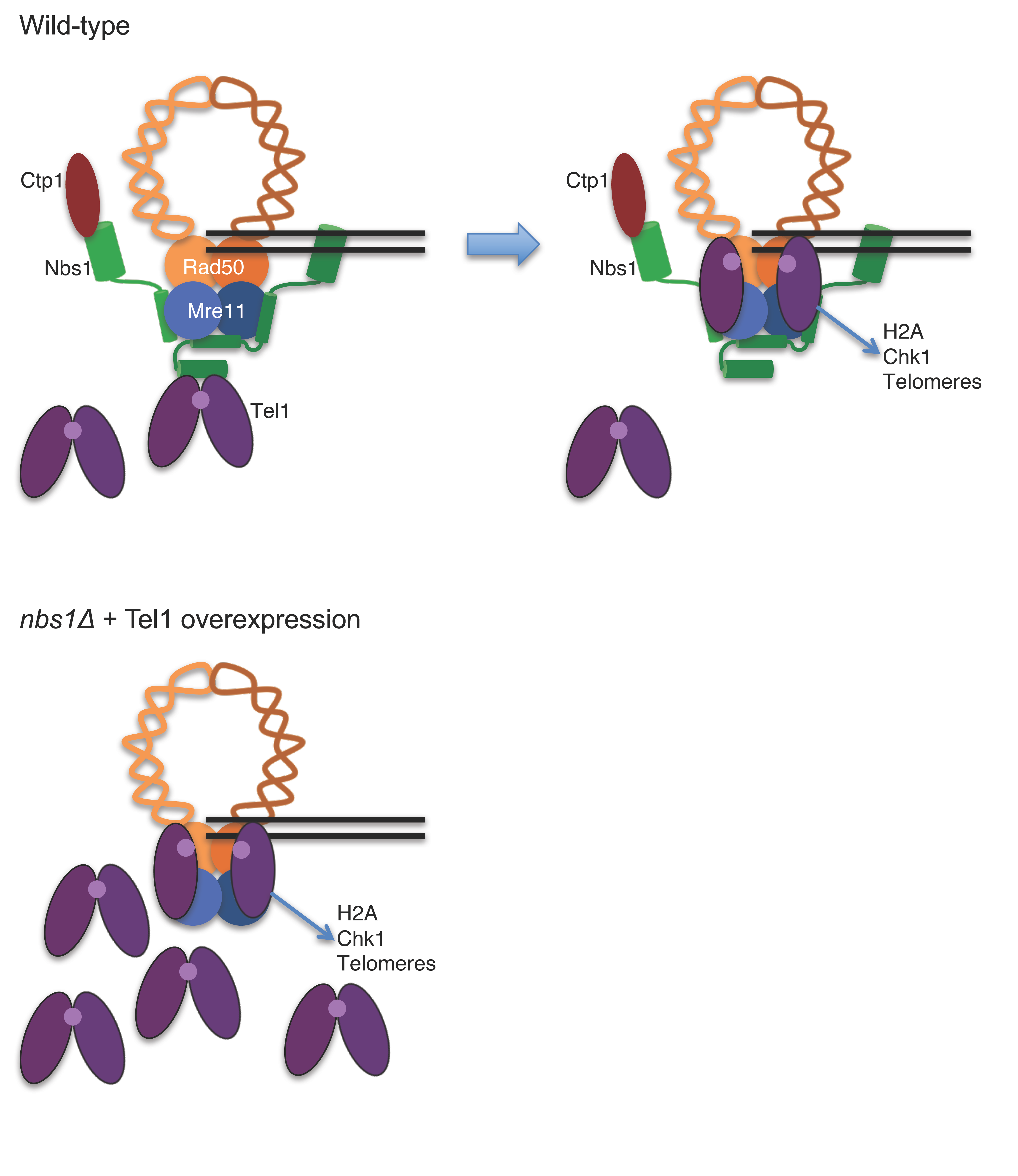
Model for MRN interactions with Tel1. In wild type, Tel1 localizes at DSB and telomeres by binding the Tel1/ATM interaction module at the C-terminus of Nbs1. This binding facilitates a lower affinity interaction with the closed conformation of Mre11-Rad50, which stimulates Tel1 activity, resulting in phosphorylation of substrates at DSBs and telomeres. Overexpression of Tel1 in *nbs1Δ* cells promotes the low affinity interaction of Tel1 with Mre11-Rad50, which partially bypasses the requirement for Nbs1 in Tel1 activity.

Mre11-Rad50 is conserved in all domains of life, whereas the Nbs1 subunit has only been identified in eukaryotes (Stracker & Petrini, 2011). In mammalian cells and budding yeast, Nbs1 is required for localization of the complex to the nucleus (Desai-Mehta et al., 2001, Tsukamoto et al., 2005). In mice, it was recently shown that only a minimal fragment of Nbs1 containing the Mre11 binding domain was required for stability of the complex, nuclear localization, DNA binding, and nuclease activities of Mre11-Rad50 (Kim et al., 2017). This fragment lacked the C-terminal ATM binding domain, yet ATM activity was not completely abolished. In budding yeast, it was recently reported that fusion of a nuclear localization signal (NLS) to Mre11 was sufficient to retain nuclease functions of the Mre11 complex independent of Xrs2 (Nbs1), but not Tel1 activity (Oh et al., 2016). Our data presented here demonstrate that *S. pombe* Nbs1 is completely dispensable for the formation of Mre11-Rad50 protein complex and its localization at DSBs. This complex is unable to catalyze DSB repair, presumably because Nbs1 is required to recruit Ctp1, which is essential for DNA end processing and resection by Mre11 complex (Lloyd et al., 2009, Williams et al., 2009). However, the meiotic defects caused by *nbs1Δ* are less severe than those caused by *mre11Δ, rad50Δ* or *ctp1Δ* (Milman et al., 2009), suggesting that Mre11-Rad50 retains a weak Ctp1-dependent DNA end processing activity in the absence of Nbs1, at least during meiosis.

The first crystallographic structure of an Mre11-Nbs1 interface showed that two monomers of Nbs1 bind the Mre11-Rad50 globular domain asymmetrically through a region at the C-terminus of Nbs1 (Schiller et al., 2012). The Tel1-binding module of Nbs1 lies immediately downstream of the Mre11 interaction region, which suggests Tel1 localizes near the Mre11-Rad50 globular domain. This architecture, along with the absolute requirement for Mre11-Rad50 in DNA damage-induced Tel1 activity, as shown here, strongly suggests that in addition to the well-established Nbs1-Tel1 interaction interface, another interface also exists between Mre11-Rad50 and Tel1. This model is supported by *in vitro* gel filtration evidence indicating that ATM has an affinity for Mre11-Rad50, as well as *in vitro* studies reporting that Mre11-Rad50 stimulates ATM-mediated p53 phosphorylation (Lee & Paull, 2004).

How Mre11-Rad50 stimulates Tel1 activity remains enigmatic. Our data shows that Mre11 endonuclease activity is dispensable for Tel1 activity, which is consistent with previous data in mice and with purified human proteins (Buis et al., 2008, Lee et al., 2013, Lee & Paull, 2005, Limbo et al., 2011). Despite the dispensability of Mre11 nuclease activity, the presence of Rad50 at DSBs alone was insufficient to elicit Tel1 activity. TAP-Rad50-I1192W, which is unable to efficiently form the closed conformation of Mre11-Rad50, was unable to stimulate Tel1 activity towards H2A in response to IR, even when Tel1 was overexpressed. The conformation of Mre11-Rad50 greatly influences ATM activity, with ATM activation occurring in the ATP-bound, closed conformation (Lee et al., 2013, Williams et al., 2011). Our data provide *in vivo* evidence that both the recruitment of Tel1 by Nbs1 and the stimulatory role of Mre11- Rad50 occur prior to ATP hydrolysis.

In the absence of DNA damage, ATM exists as an inactive homodimer. Exposure to ionizing radiation induces monomerization, which exposes the kinase domain and allows ATM to phosphorylate its substrates. In human cells, ATM monomerization is catalyzed by autophosphorylation at the serine residue at position 1981 (Bakkenist & Kastan, 2003). However, the importance of this autophosphorylation is controversial, as mutation of the homologous residue in murine ATM (S1987) does not significantly impair ATM activity (Daniel et al., 2008, Pellegrini et al., 2006). Cryo-EM structures of *S. pombe* Tel1 homodimers demonstrated that this serine residue lies in a 32-amino acid insertion (termed INS32) that is absent in *S. pombe* Tel1 (Wang et al., 2016). As with murine ATM (Pellegrini et al., 2006), autophosphorylation may be unnecessary for Tel1 activation in *S. pombe.* However, as seen with other members of the PIKK family, its activity may be inhibited through the blockage of the kinase domain of one molecule of Tel1 with another. Thus, the most conserved property of ATM/Tel1 activation appears to be the disengagement of the homodimer. It is tempting to speculate that Mre11-Rad50 mediates ATM/Te1l activity through ATM/Te1l monomerization.

Although we failed to detect an IR-induced increase in γH2A formation when Tel1 was overexpressed in *mre11Δ rad3Δ* or *rad50-I1192W rad3Δ* backgrounds, the untreated samples had a significant increase in basal γH2A levels when compared to the empty vector controls (Figs 3A and 7B). It remains to be determined whether this γH2A formation occurs randomly in chromatin or in response to specific events such as replication fork collapse or telomere erosion. Whichever is the case, it is evident that Tel1 overexpression restores substantial Tel1 activity even in the absence of MRN complex.

Oxidative stress was reported to cause MRN-independent ATM activation by a pathway that does not involve DNA damage (Guo et al., 2010). Interestingly, unlike the MRN-dependent pathway of human ATM activation, in which an inactive ATM dimer disengages into active monomers after autophosphorylation, ATM activation from oxidative stress exists as a disulfide-linked covalent dimer formed through the C-terminus of ATM. Mutation of this C-terminal region specifically abolished ATM activation caused by oxidative stress but retained activation stimulated by DNA damage. Moreover, Nbs1-independent ATM activation has been observed by assaying phosphorylation of p53 in postmitotic neural tissue (Frappart et al., 2005, Li et al., 2012). Thus, MRN is not absolutely required for all ATM activity, but appears to be critical in the context of DSBs and telomeres.

When ATM binds MRN at DSBs, it phosphorylates histone H2AX in surrounding chromatin, which then binds the C-terminal BRCT domains of the DNA damage mediator protein, MDC1. MDC1 binds the FHA/BRCT domains of NBS1 and autophosphorylated ATM, which increases ATM signaling at DSBs (Lou et al., 2006, Stucki et al., 2005). *S. pombe* has an MDC1-like protein known as Mdb1, which binds γH2A through its C-terminal BRCT domains (Wei et al., 2014). An FHA-like structure at the N-terminus of Mdb1 mediates homodimerization, analogous to MDC1 (Luo et al., 2015). Mdb1 might mediate the MRN-independent activity of Tel1 reported here.

In summary, our study underscores the importance of MRN, and particularly Nbs1, in Tel1 activity at DSBs and telomeres. In addition, we provide *in vivo* evidence of a critical stimulatory role of Mre11-Rad50 in the Tel1 signaling pathway. We propose that recruitment of Tel1 to DNA ends is principally dependent on high affinity binding to Nbs1, whereas activation of Tel1 after it binds Nbs1 involves a lower affinity interaction with Mre11-Rad50. This sequential mechanism of recruitment and activation of ATM/Tel1 may play an important role in coordinating its activity with DSB repair and telomere maintenance.

## Materials and Methods

General *S. pombe* methods used have been previously described (Forsburg & Rhind, 2006). Strains used are listed in Supplementary Table S1. For DNA damage sensitivity assays, 5-fold serial dilutions of log-phase cells were spotted onto agar plates and treated with the indicated dose of DNA damage. Chromatin immunoprecipitation experiments were performed as previously described (Limbo et al., 2007) and are representative of at least 2 independent experiments. HO-endonuclease expression was driven from the thiamine repressible *nmt41* promoter. Samples were taken at indicated time points after removal of thiamine. The Mre11-TBD fusion construct was generated by amplifying the 3’ end of *nbs1*^+^ using primers containing homologous regions to 3’ end of *mre11*+. The PCR product was transformed into wild-type cells and checked for proper integration. Details and primers sequences are available upon request.

Western blots and co-immunoprecipitation experiments were performed as previously described (Limbo et al., 2012). Experiments were done with asynchronous cells grown to log-phase. Where indicated, cells were treated with 90 Gy of ionizing radiation from a Cs-137 source and harvested 30 minutes after exposure. Membranes were blotted with one of the following antibodies: PAP (Sigma P1291), MYC (Covance MMS-150P), FLAG (Sigma F3165), Tubulin (Sigma T5168), HA (Roche 11666606001), and total H2A (Active Motif 39235). The anti-γH2A antibody was previously described (Rogakou et al., 1999).

For telomere Southern blots, *mre11Δ* or *nbs1Δ* strains were crossed to *rad3Δ* with *tel1*^+^ either under its endogenous promoter or the full-strength thiamine-repressible *nmt1* promoter. Confirmed strains were then streaked for single colonies sequentially, with a liquid culture grown at each passage for genomic DNA extraction. Strains were grown and maintained on minimal media lacking thiamine to ensure full expression of *tel1*^+^ for the duration of the experiment. For the Mre11-TBD fusion Southern blot, generated mutants were streaked 10 times sequentially prior to isolation of genomic DNA to allow for circularization of chromosomes. Southern blotting was performed as previously described (Limbo et al., 2012). Briefly, DNA was digested with *ECo*RI and resolved on 2% TAE agarose gels. DNA was transferred to a nylon membrane by capillary method and incubated with TAS1 probe (Nakamura et al., 1998) generated by PCR with biotinylated dCTP. The membrane was incubated with dye-labeled streptavidin and scanned on a LI-COR Odyssey imaging system. An alternate pathway of Tel1 recruitment to telomeres independently of the C-terminus of Nbs1 has been previously described (Subramanian & Nakamura, 2010). In this alternate pathway, Tel1-mediated telomere maintenance was observed in a pathway that depended on the N-terminus of both Rad3 and Nbs1. To exclude this alternate pathway, we used full deletions of both of these proteins in both our DNA damage and telomere assays. Moreover, this alternate pathway appears to be specific to telomeres (Fig EV2).

## Acknowledgements

We thank Christophe Redon for providing the γH2A antibody and Michael Nick Boddy, John Tainer, and members of the Russell Lab for invaluable discussions. This study was supported by a fellowship from The Uehara Memorial Foundation awarded to YY and National Institutes of Health grants GM059447, CA077325 and CA117638 awarded to PR.

## Author Contributions

OL, YY, and PR designed experiments and analyzed results. OL and YY performed experiments. OL and PR prepared manuscript.

---

**Figure EV1.**
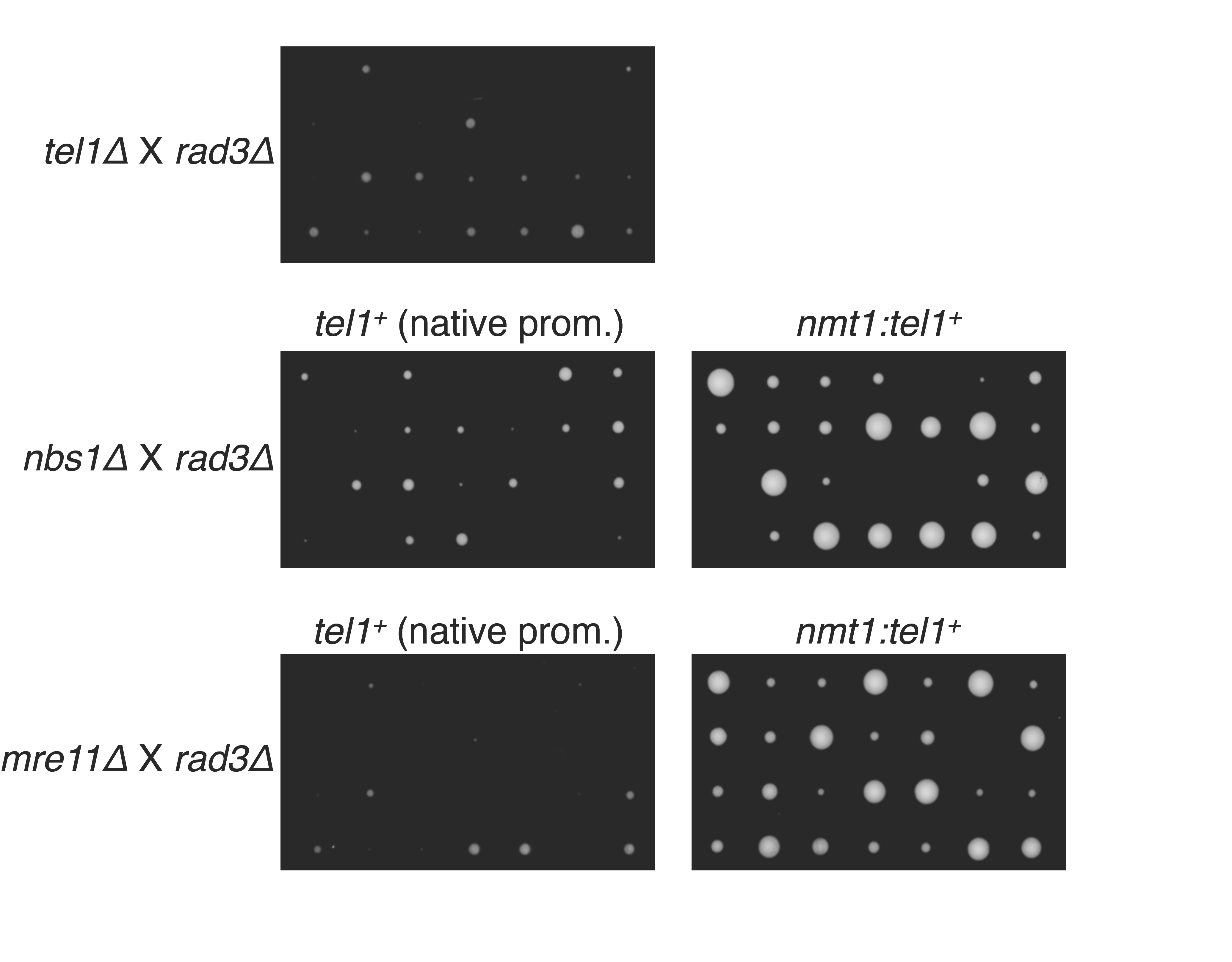
Cells overexpressing Tel1 grow faster than cells with endogenous levels. Cells with the indicated genotype were crossed with either *tel1*^+^ under its native or the *nmt1* overexpression promoter. Tetrad dissection was performed on the same day with pictures taken after 4 days of growth at 30°C

**Figure EV2.**
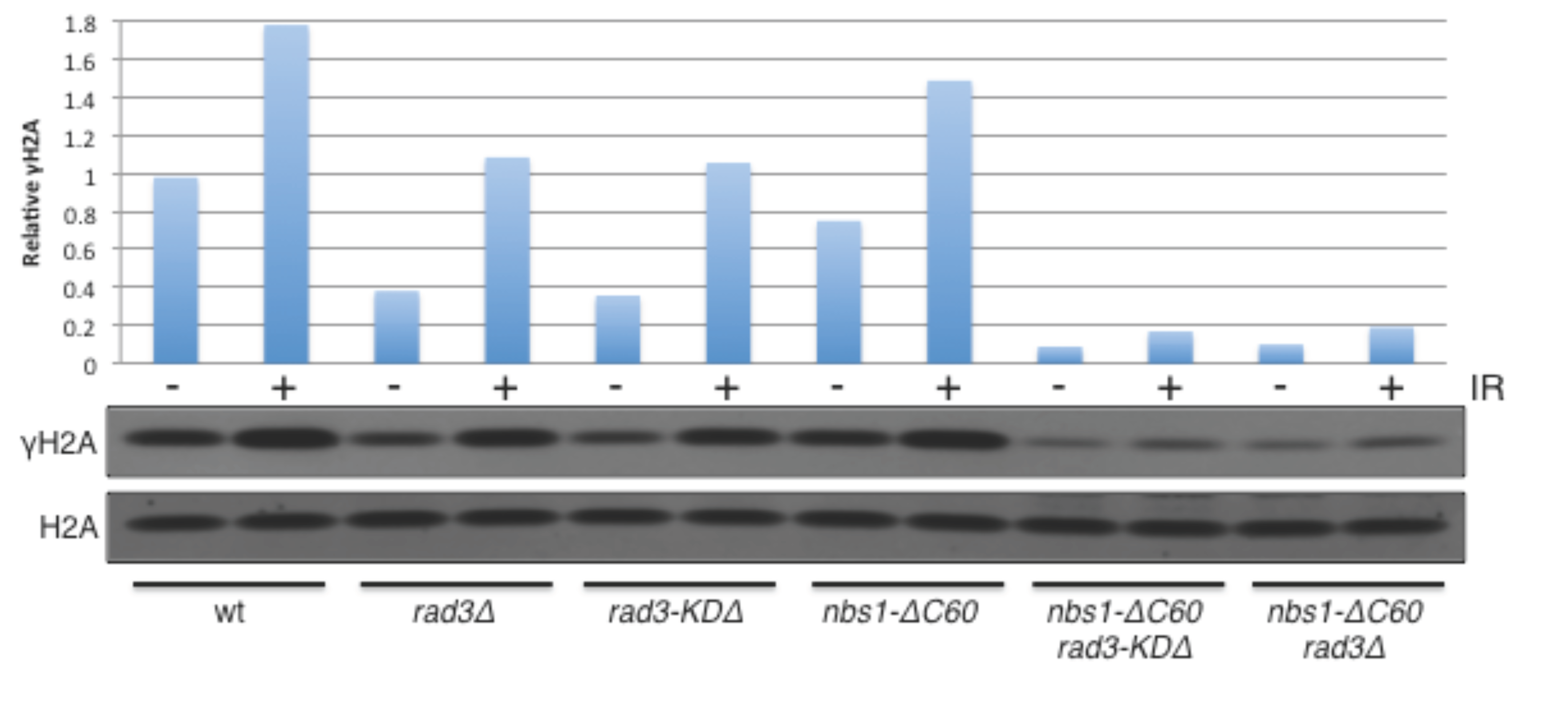
The alternative method of Tel1 recruitment to telomeres dependent on the N-terminus of both Rad3 and Nbs1 is not sufficient for Tel1 activity towards γH2A in response to ionizing radiation Rad3-KDΔ truncates the C-terminus of Rad3 containing the kinase domain. The *nbs1-ΔC60 rad3-KDΔ* strain was previously shown to be sufficient for Tel1 activity at telomeres.

